# Filter paper-based spin column for low throughput nucleic acid purification

**DOI:** 10.1101/392696

**Authors:** Rui Shi, Ramsey S. Lewis, Dilip R. Panthee

## Abstract

We describe a method of recharging used spin column or assembling homemade spin column using filter paper as binding material for low throughput nucleic acid purification. We evaluated the efficiency of filter paper based spin columns in the purification of different type of nucleic acids. For instance, by following protocols of respective commercial kits, we found that filter paper to be a useful binding material for purification of many types of nucleic acids, including plant genomic DNA, plant total RNA, PCR product, and DNA from agarose gels. We also found that filter paper has a weak binding affinity to plasmid DNA in tested miniprep protocols. Also, we present the protocols of using filter paper recharged spin column or homemade spin column for low throughput purification of plant genomic DNA and plant total RNA with commercial kit buffer leftover and less expensive homemade buffer.

## Introduction

As a basic molecular biology technique, nucleic acid purification is the starting point for many molecular biology applications [1, 2]. Classic nucleic acid purification methods are based on organic extraction followed by ethanol-based precipitation. However, classic methods are time-consuming and require the use of toxic solvents, such as chloroform and phenol which can be harmful to the user and the environment [1, 2].

Commercial kits usually follow solid-phase purification approaches whereby nucleic acids in an extraction solution are absorbed by a solid-phase binding material under appropriate chaotropic conditions, followed by washing of non-nucleic acids remaining on the binding material using appropriate buffer solutions, and elution of purified nucleic acids from the binding material using low salt solutions [1, 3, 4]. Commercial kits using this approach permit for fast purification of high-quality nucleic acids, although they are expensive to use.

The success of commercial kits largely relies on the spin columns or spin plates assembled with solid-phase nucleic acid binding material which allow easy binding, washing, and elution of nucleic acids in the purification process. The most widely adopted nucleic acid binding material in the past has been mineral based silica material in the form of a matrix (glass fiber filter or silica membrane) or powder (glass milk or silica slurry). Guanidine based buffer is usually adopted to provide a chaotropic condition for nucleic acid binding [1]. Plant-based cellulose material has been applied in DNA and RNA purification. For example, Su and Comeau [5] adopted cellulose fiber to purify nucleic acids by using sodium chloride and polyethylene glycol as binding reagents. MeganCel paramagnetic cellulose particles are an example of a commercialized product from Promega [6]. Also, modified cellulose such as DEAE-modified cellulose membranes can be applied as anion exchange materials for nucleic acid purification [7]. In a previous report, we found that filter paper made from cellulose fiber to be a viable replacement for silicon-based material for purification of plant genomic DNA from CTAB/NaCl extractions [8]. Another recent report also indicated that the filter paper tip could be used to purify nucleic acids from crude extract with NaCl in a concise time for subsequent PCR based analyses [9]. Since cellulose-based filter paper is an inexpensive and readily available material in laboratories, we hypothesized that filter paper might have more widespread usefulness in solid-phase based nucleic acid purification. We, therefore, described our attempts to use filter paper in assembling recharged or homemade spin column suitable for low throughput nucleic acids purification.

## Material and methods

### Plant materials, plasmid DNA and PCR product

Tomato (*Solanum lycopersicum L*.) plant samples were collected from the field or greenhouse of Mountain Horticultural Crops Research & Extension Center of North Carolina State University at Mills River, NC. USA. Tobacco (*Nicotiana tabacum L*.) plant samples were collected from laboratory or growth chamber experiments being carried out by the Department of Crop and Soil Science at North Carolina State University, Raleigh. For DNA purification, collected fresh samples can be used immediately, or stored at −20°C before use. Usually, about 50 to 100 mg leaf samples were collected and put into 2 ml screw cap tubes for grinding using mechanic homogenizer, or 1.5 ml Eppendorf tube for grinding using plastic pellet pestles. The sample for RNA purification should be frozen in liquid nitrogen quickly after collection and grind into fine powder in liquid nitrogen, then transfer 50 to 100 mg ground sample into the tube and to use immediately or store at −80°C before use.

pUC19 and pBI121 plasmids were purchased from Invitrogen (Carlsbad, CA). The GUS gene fragment was amplified from the pBI121 plasmid or transgenic tobacco by PCR using the following primer:

The forward primer, 5’- TGACCTCGAGGTCGACGATATCGTCGTCATGAAGATGCGGAC-3’

Reverse primer, 5’- CTAGACTAGTCCCGGGGGTACCATCCACGCCGTATTCGGTG-3’.

### Filter paper-based spin column preparation

Recharging of used commercial spin columns is initiated by separating plastic parts and treating them with 10% bleach for at least 10 mins, followed by thorough rinsing with sterilized water several times, and air-drying. Filter paper discs were punched from sheet of Whatman^™^ qualitative filter paper, Grade 3 (GE Healthcare Life Science, UK) or equivalent filter paper using 3/16-inch (~8mm) paper puncher, and then one or two layers of filter paper disc is added to the column by pushing down them to bottom of column tightly using the end of a 200 μl pipette tip (Fig 1A). The assembled spin column could be autoclaved and air dried.

**Fig 1.**
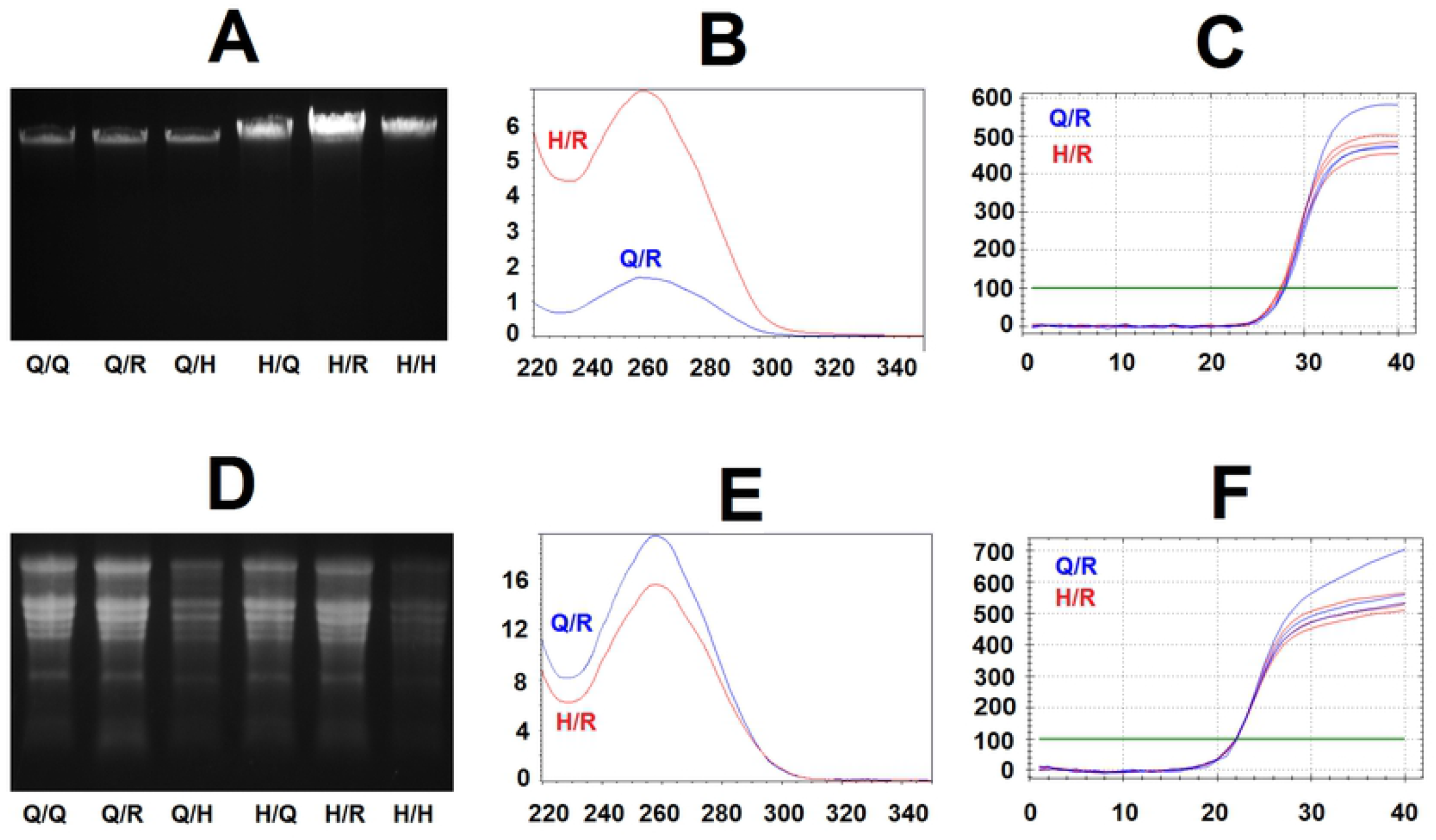
Illustration of recharging spin column and homemade spin column using filter paper. (A) Recharged used spin column with a flat bottom and net structure to support filter paper discs. (B) Homemade filter paper-based spin column prepared by using 0.5 ml tube with the bottom part cut, and adding the top part of 10 μl pipette tip as supporting tube to support two layers of filter paper discs loaded on it within the tube. The homemade spin column is based on 0.5 ml PCR tubes. In brief, the bottom of 0.5 ml tubes is initially cut off, followed by the insertion of the upper part of 10 μl pipette tips to serve as supporting tubing, then load one or two layers of filter paper discs on the supporting ring in tube, and push tightly using upper end of 10 μl tip (Fig 1B). For homemade spin column based on 0.5 ml PCR tube, filter paper discs were punched using a standard 1/4-inch (6.35 mm) paper puncher from filter paper sheet.

### Purify plant genomic DNA using filter paper-based spin column with Qiagen plant DNeasy kit buffer or homemade buffer

This protocol is an example of using filter paper-based spin columns to purify plant genomic DNA with Qiagen DNeasy Plant mini kit buffers referred to Qiagen DNeasy^®^ plant handbook (March 2018) or less expensive homemade buffer as described by Lemke et al [10].

To lysis plant material, add 400 μl AP1 buffer of Qiagen kit or homemade lysis buffer (0.5% SDS, 8% PVP-10, 250 mM NaCl, 25 mM Na-DETA, 200 mM Tris-HCl pH7.5) along with 4 μl RNase A stock solution (100 mg/ml, Qiagen) into 2 ml screw cap tube with 50 to 100 mg frozen or fresh plant material, then add two tungsten carbide beads (Qiagen) or similar beads into tube and homogenize samples using homogenization equipment like FastPrep FP120 cell homogenizer (Savant Instruments Inc, Holbrook, NY) followed the manual. Alternatively, grind 50 to 100 mg frozen or fresh sample in 1.5 ml Eppendorf tube with 400 μl lysis buffer and 4 ul RNase A stock solution using disposable plastic pellet pestles (USA Scientific, Ocala, FL).

Tubes with lysis mixture are incubated at 65°C for 10 min (mix by inverting the tube 2 to 3 times), then add 130 μl P3 buffer of Qiagen kit or 130 μl homemade precipitation buffer (5M KAc, pH 6.5) and maintenance on ice for at least 5 mins. Centrifuge sample tube using a microcentrifuge at its top speed (≥16,000 g) for 10 mins to clear lysate.

In case lysate is not well cleared, an option step followed is to place a spin column with one layer of filter paper into a new 2 ml tube which is used as collection tube, then transfer lysate to the spin column and centrifuge at 6,000 × g for 1 min followed another round of centrifuge at top speed for 1 min to get cleared lysate in collection tube.

To bind DNA onto filter paper discs in spin column, transfer cleared lysate from collection tube into a new tube and mix with 1.5 volume of AW1 buffer of Qiagen kit or homemade binding buffer (2M Guanidine Hydrochloride, 75% ethanol) by pipetting or inversion, transfer mixture to new spin column assembled with two layers filter paper disc, which placed in a new 2-ml collection tube, centrifuge at 6000 × g for 1 min, discard flow through and repeat to load remaining solution for another round of centrifuge step to allow all cleared lysate flow through filter paper discs in spin column.

To wash filter paper discs in the spin column, add 500 μl AW2 buffer of Qiagen kit or homemade washing buffer I (10 mM NaCl, 10 mM Tris-HCl pH6.5, 80% ethanol) to the column, spin at 6000 × g 1 min, discard flow through and reuse collection tube. Add 500 μl AW2 buffer of Qiagen kit or homemade washing buffer II (95% ethanol), spin at 6000 g for 1 min and then transfer spin column to the new collection tube. Centrifuge columns at the top speed for at least 2 mins.

For DNA elution from filter paper discs, insert a spin column to new 1.5 ml collection tube, air dry for a while and add 100 μl AE buffer of Qiagen kit or 10 mM Tris-HCl (pH8.5) on filter paper in spin column, and maintain at room temperature for 5 mins, then centrifuge at 6000 g for 1 min to elute DNA into collection tube.

### Purification of plant RNA using filter paper-based spin column with commercial kit buffer or homemade buffer

This protocol is an example of using filter paper-based spin columns to purify plant total RNA with Qiagen RNeasy Plant mini kit buffers according to RNeasy^®^ Mini Handbook (Fourth edition, June 2012) or less expensive homemade buffer presented by Yaffe et al [11].

In brief, add up to 100 mg grinded frozen sample into 1.5 or 2 ml tube, add 450 μl buffer RTL or RLC of Qiagen kit with 1% (v/v) b-mercaptoethanol or homemade lysis buffer (8 M guanidine hydrochloride, 20 mM MES hydrate and 20 mM EDTA) along with 1% (v/v) b-mercaptoethanol.

Tube with lysis mixture is incubated at 55°C for 1 to 3 min (mix by inverting the tube 2 to 3 times), and then centrifuge tube using a microcentrifuge at top speed (≥16,000 × g) for 5 to 10 mins to clear lysate. If lysate is not clear enough, then insert a spin column assembled with one layer of filter paper in a 2 ml new collection tube, and transfer all cleared lysate to spin column and centrifuge at 8,000 × g for 1 min followed another round of spin at top speed for 1 mins to get cleared lysate.

Transfer all cleared lysate to a new 1.5 ml tube and mix with half amount of ethanol by pipetting or inversion. Place a new spin column assembled with two layers of filter paper discs into a new 2 ml collection tube, then transfer lysate/ethanol mixture to spin column and centrifuge at 8000 × g for 1 min, discard flow through, and reuse collection tube and repeat to load remaining solution and centrifuge again.

To wash column, add 700 μl RW1 buffer of Qiagen kit or homemade washing buffer I (3 M Na-acetate pH 5.2) to the spin column, then spin at 6000 × g 1 min and discard flow through from collection tube. Add 500 μl RPE buffer of Qiagen kit or homemade washing buffer II (70% ethanol), and centrifuge at 8000 × g for 1 min, transfer spin column to new collection tube and centrifuge spin column at the top speed for at least 2 mins to eliminate remaining ethanol. Insert a spin column to new 1.5 ml collection tube, air dry for a while, then add 50 μl RNase free water and maintain at room temperature for 1 mins, then centrifuge at 8000 × g for 1 min to elute RNA.

For RNA purification, all reagent and buffer should be treated by diethyl pyrocarbonate (DEPC) to eliminate RNase and also keep working area clean.

### Evaluation of purified nucleic acids

DNA yield and quality were checked by 1 to 1.5 % agarose electrophoresis stained using Ethidium Bromide (EB), RNA integrity and quality were evaluated by 1 % MOPs-formaldehyde denaturing agarose electrophoresis with EB in RNA loading buffer [12]. Ethidium bromide stained DNA, or RNA fluorescent bands were visualized and recorded by Bio-Rad Gel Doc^™^ XR+ gel image system (Bio-Rad Laboratories, Hercules, CA), or FOTODYNE system (FOTODYNE incorporated, Hartland, WI).

DNA and RNA quality and quantity were also estimated by Nanodrop 2000 UV spectrometer (Fisher Scientific, Waltham, MA). Also, DNA concentration was measured using Hoefer fluorometer DQ300 (Hoefer, Holiston, MA) with double DNA specific H33258 dye (Sigma-Aldrich).

Quantitative PCR (qPCR) and quantitative reverse transcription PCR (qRT-PCR) approaches were used to further precisely check whether the purified DNAs and RNAs contain contamination which could interfere with the efficiency of PCR reaction. In brief, 10 μl SYBR green-based qPCR reaction was prepared using Luna^®^ Universal Probe qPCR Master Mix (New England Biolabs, Ipswich, MA, USA) with 0.5 μM primer set NtTublin_1 for tobacco [13]. For DNA evaluation, we added 50 ng genomic DNA purified by different approaches in reaction. For RNA evaluation, cDNAs were reverse transcribed from 200 ng purified RNA in 10 μl reverse transcription reaction prepared using iScript^™^ cDNA Synthesis Kit (Bio-Rad), and cDNA amount to 2.5 ng RNA was added to the qRT-PCR reaction. Both qPCR and qRT-PCR were run on CFX96^™^ Real-Time System followed standard two step PCR program as suggested by Luna^®^ Universal Probe qPCR Master Mix manual. Amplification efficiencies of different input templates were evaluated using CFX Maestro^™^ Software version 1.1 (Bio-Rad) based on quantification cycle (Cq) value [14], which is calculated fractional cycle at which the target DNA amplicon associated fluorescent accumulated up to an arbitrary threshold.

## Results

### Recharged used spin column and homemade spin column using filter paper

We previously described a method to prepare filter paper-based 96 well spin plates suitable for high throughput nucleic acid purification [8], and another recent publication also reported filter paper dipstick designed for fast purification of DNA and RNA used in PCR based applications that do not require large quantities of nucleic acids [9]. However, compared to the multiple well spin plate and dipstick, spin column assembled with filter paper should be the more appropriate format for low throughput nucleic purification conducted in most biological labs. Therefore, we have made attempts to prepare a filter paper based spin column.

Our efforts start from recharging of used commercial spin columns with filter paper disc(s). As described in the Methods section, such recharged spin column can be prepared based one a commercial spin column which has a flat bottom but equipped with net structure to support binding material, such as those Wizard^®^ SV minicolumns of Promega (Madison, WI) (Fig 1A). Alternatively, a spin column with a conical bottom (V shape bottom) equipped drip opening, such as a miniprep column of Qiagen can be adopted. Both formats are convenient for reloading of filter paper discs with a diameter of 5/16 inch (~8 mm). A recent version of a spin column named as a microspin column, such as those used in NEB kit can be recharged by using filter paper discs with a diameter of 5/16 inch or using filter paper disc(s) with a diameter of 3/16 inch (6.35mmm).

Since filter paper is relatively physically stronger to hold its shape compare to the soft silica based membrane, such as glass fiber filter, therefore, we prefer to exclude fixing ring (O-ring) and support plastic frit adopted in many types of the commercial spin column in the recharging of these used spin column. The advantage of excluding these items is to simplify the recharging process, and also help to avoid the problem of solution leftover on the fixing ring during purification experiment.

In addition to the recharged used spin column, we also try to prepare homemade spin column assembled with filter paper. In the previous report, Borodina et al [15] have described a way to prepare glass fiber filter based homemade spin column using a 0.5 ml PCR tube. They added silica based glass fiber material to the bottom of the tube with several small holes punched, however, it is harder to push filter paper discs down to the bottom and seal the bottom of the tube properly like soft glass fiber filter adopted in the homemade silica spin column. For example, if we push filter paper too hard, then filter paper did fill the bottom of the tube but might be too tight to block flow through of solution. Otherwise, it might leave space and result in leaking of the sample solution.

After many attempts, we found that adding upper part of 10 ul tip as supporting ring in the tube, and then can load filter paper disc(s) on the supporting ring to form a simple homemade spin column. For better performance, we chose the upper part of the tip with additional edges (Fig 1B), which can provide enough supporting area to support filter paper disc and also leave space between supporting ring and wall of the tube for allowing solution flow through filter discs without leftover.

### Evaluation of filter paper in the purification of different nucleic acids

Though filter paper has been successfully adopted in method using NaCl as a binding reagent [5, 8, 9], we also found that filter paper work well in DNA purification followed protocol using guanidine-based reagent, which is commonly adopted in commercial kit for nucleic acid binding, and therefore, it is possible to use filter paper-based spin column as substitute for nucleic acid purification using commercial kit.

To investigate whether filter paper-based spin column can be used in commercial kit, we reassembled spin columns of Qiagen kit by just replacing of original silicon membrane with filter paper discs, then apply such reassembled spin column in purification of different types of nucleic acids using respective Qiagen kits, including DNeasy^®^ plant mini kit for plant genomic DNA, RNeasy^®^ plant mini kit for plant total RNA, QIAquick PCR purification kit for PCR product, QIAquick gel extraction kit for DNA in agarose gel, and QIAprep spin miniprep kit for plasmid DNA. For comparison of effectiveness, we included original spin columns and reassembled spin column using glass fiber filters (Whatman^™^ glass microfiber filters, Grade GF/F, GE), a silica based material usually adopted for recharging or preparing homemade spin column [10, 15].

In these experiments, the filter paper based-spin columns shown better performance for purification of tomato genomic DNA and yield higher amount final DNA than two silica-based spin column, including original Qiagen kit spin column and glass fiber-reassembled spin column, (Fig 2A). Filter paper-based spin column was also functioned in purify plant total RNA (Fig 2B), DNA from PCR (Fig 2C) and recover DNA from agarose gel (Fig 2D), although the yields of final nucleic acid are relatively lower (half) compare to yield from experiments using silica-based spin columns.

**Fig 2.**
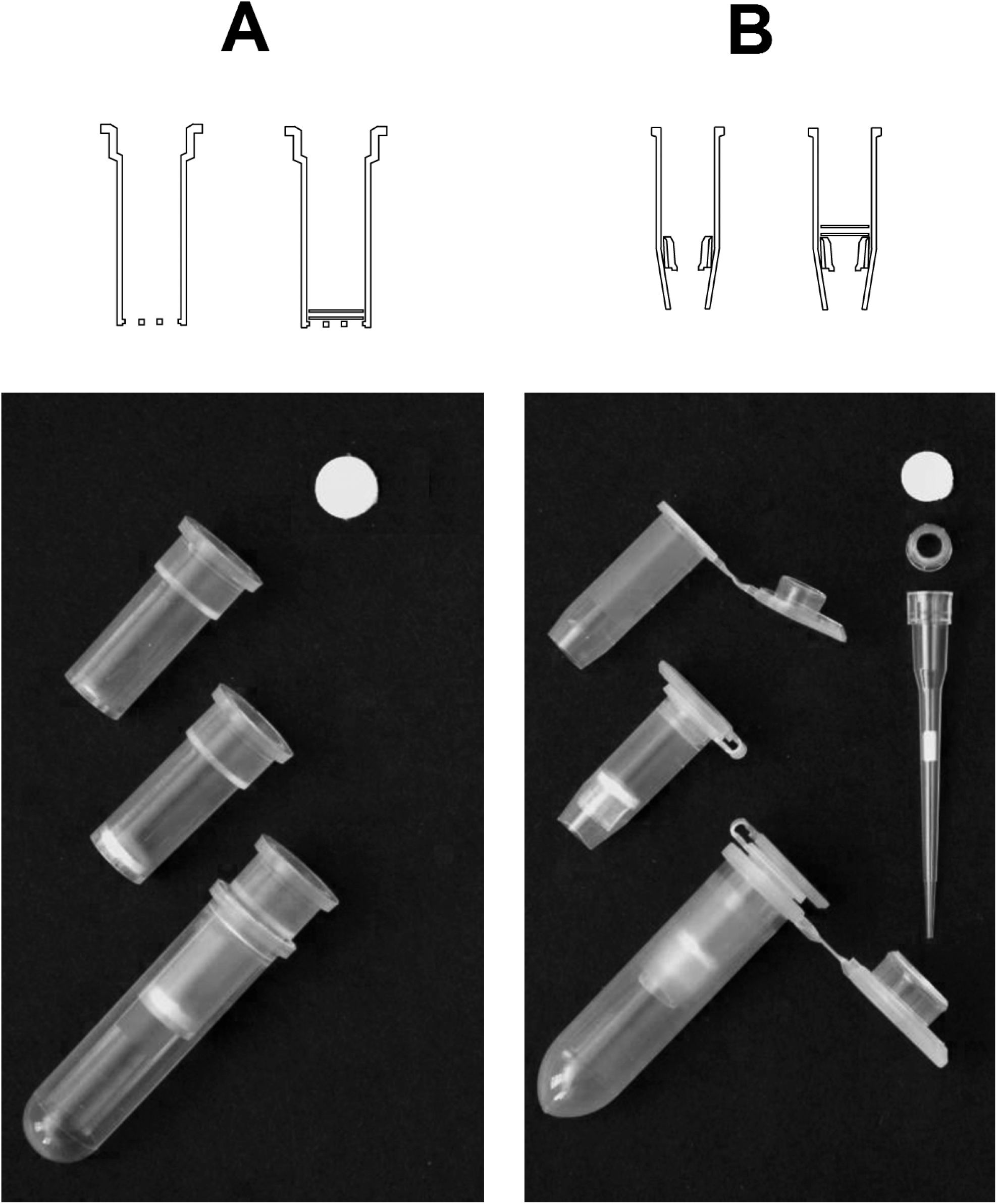
The efficiency of filter paper for purification of different types of nucleic acid using respective Qiagen kits. (A) Tomato genomic DNAs purified using Qiagen DNeasy^®^ plant mini kit. (B) Tomato total RNAs purified using Qiagen RNeasy^®^ plant mini kit. (C) PCR products of GUS fragment purified Qiagen QIAquick PCR purification kit. (D) PCR products of GUS fragment recovered from agarose gel using a Qiagen QIAquick gel extraction kit. (E) pUC-19 plasmid DNAs purified Qiagen QIAprep spin miniprep kit.

For each panel, lane from left to right labeled with original column, glass fiber and filter paper is the same volume of nucleic acid elution from purification experiments using original Qiagen spin column, reassembled spin column using two layers of Whatman^™^ glass microfiber filters, (Grade GF/F), and reassembled spin column using two layers of Whatman^™^ qualitative filter paper, (Grade 3) respectively.

Apart from these finding, we also noticed that filter paper reassembled spin column does not work well for purification of plasmid DNA using QIAprep spin miniprep kit protocol since only a weak plasmid DNA band was detected by agarose gel electrophoresis (Fig 2E).

According to these results, it seems that filter paper works well for purifying long linear double strand nucleic acids, such as plant genomic DNA followed protocol for silica-based binding material. On another hand, the filter paper was substantially less effective for supercoiled plasmid DNAs in the present experiments. Nevertheless, these experiments still indicated that filter paper could serve as an alternative binding material to replace silicon material for purification of many types of nucleic acids followed the protocols for silica-based material, except plasmid DNA using protocols developed for silica based nucleic binding material.

### Purification of nucleic acids using filter paper-based spin column with commercial kit buffer or homemade buffer

Since it was confirmed that filter paper-based spin column could apply in the purification of many types of nucleic acids followed the protocol of commercial kit originally optimized for the silica-based spin column, we expected that filter paper based spin column can use as a substitute of the commercial spin column in an experiment using the commercial kit buffer leftover to save resources. Also, filter paper-based spin column might also work for a homemade buffer which mimics commercial kit’s for further reduce the expanse in the laboratory. For instance, as to our interested plant nucleic acids, there are already several in-house protocols developed for purification using the silica based material for purification of plant DNA [10, 16–18] or plant RNA [11, 19]. Therefore, we have tried this idea using tobacco plant material.

Followed the protocol described in material and method, we have successfully purified tobacco genomic DNA using filter paper-based recharged or homemade spin column following protocol using Qiagen kit buffers and an in-house protocol modified from Lemke et al [10] using homemade buffers. As shown in Fig 3A, agarose gel electrophoresis can clearly show the similar performance of purification of tobacco genomic DNA using Qiagen original spin column; filter paper recharged spin column and filter paper based homemade spin column with a buffer of Qiagen. The EB stain bands of genomic DNA band purified from the tobacco leaf by using either Qiagen original spin column, filter paper recharged spin column, and homemade spin column with Qiagen kit buffers are similar in size (integrity) and density (yield), i.e., the filter paper based spin column can be substitute of commercial silica based spin column in plant DNA purification using commercial kit buffer. On another hand, our results also indicated the higher yield of genomic DNAs purified from tobacco sample using homemade buffer compared to experiments using Qiagen kit buffer, especially for filter paper recharged spin column.

**Fig 3.**
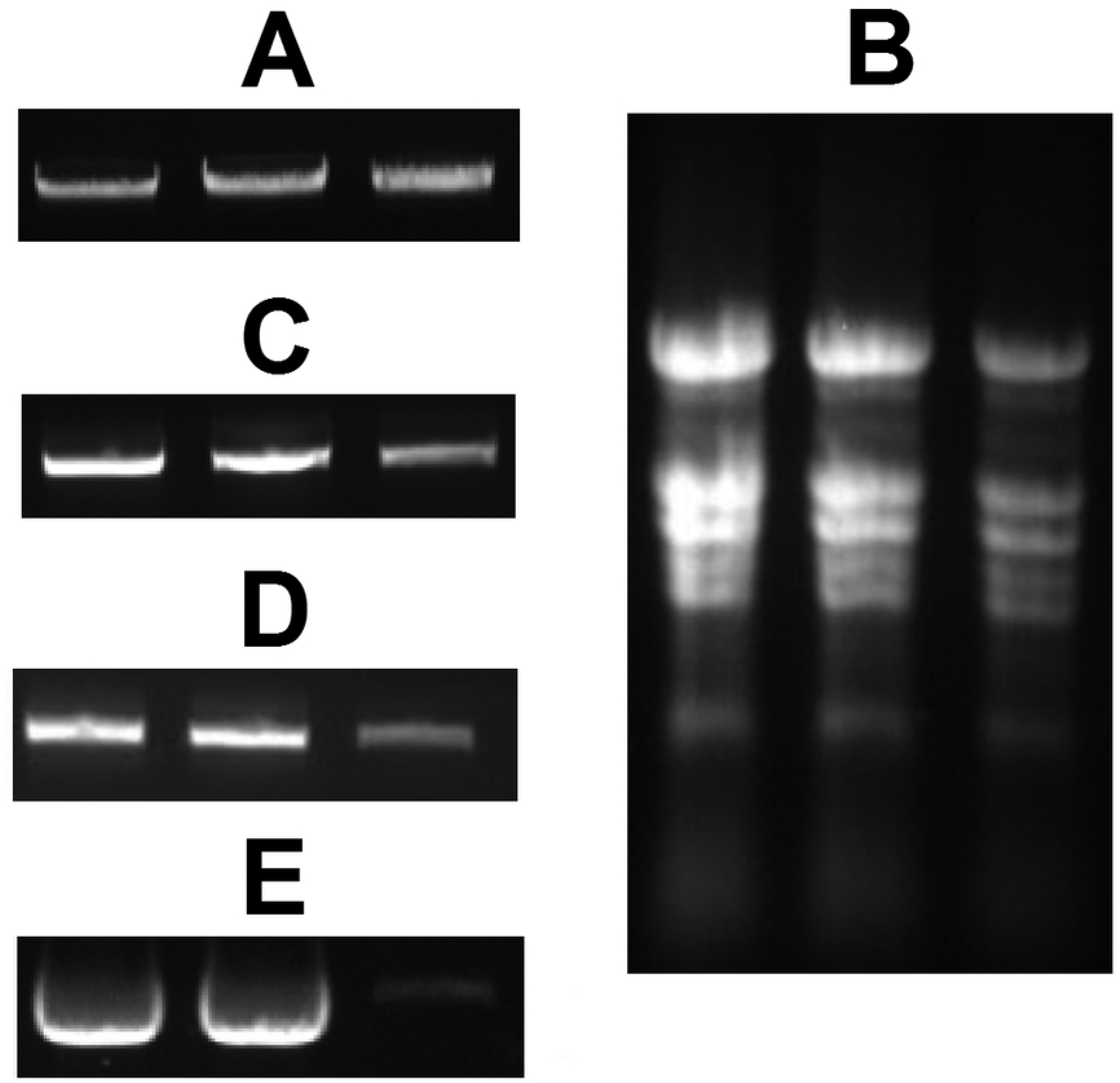
Evaluation of purification of tobacco genomic DNA and total RNA using filter paper-based spin column with respective Qiagen kit buffer and homemade buffer. (A) Agarose gel electrophoresis for 2.5 μl tobacco genomic DNAs elution from purification experiments using Qiagen DNeasy^®^ plant mini kit buffers with Qiagen original spin column (Lane Q/Q), filter paper recharged used spin column (Lane Q/R) and filter paper-based homemade spin column (Lane Q/H), followed by tobacco genomic DNAs purified using homemade buffer with Qiagen original spin column (Lane H/Q), filter paper recharged used spin column (Lane H/R) and filter paper-based homemade spin column (Lane H/H). The start material amount is 100 mg tobacco young leave tissue for experiments using a Qiagen spin column or filter paper recharged spin column, and 50 mg plant sample for homemade spin column purification. Final DNAs were eluted using 100 μl elution buffer in these experiments. (B) Typical UV spectrum curve of tobacco DNAs purified using filter paper recharged spin column with Qiagen kit buffer (Q/R, blue curve) or homemade buffers (H/R, red curve) from the same amount leaf tissue. Y-axis is UV absorbance, and X-axis is wavelength (nM). (C) Amplification plots for three duplicated qPCR reactions contain 50 ng DNA purified using Qiagen kit (Q/Q, Blue curves) and three qPCR reaction duplicates for 50 ng DNA purified from filter paper recharged spin column with homemade buffer (H/R, Red curves). The x-axis is PCR cycle numbers, Y-axis is level of SYBR fluorescence, and the green line is an arbitrary threshold to determine the Cq value (the fractional cycle number at which amplification curve meet threshold level). (D) MOPS-formaldehyde denaturing agarose gel electrophoresis separated 5 μl RNA elution from purification experiment using Qiagen RNeasy^®^ plant mini kit buffers with Qiagen original spin column (Lane Q/Q), filter paper recharged used spin column (Lane Q/R) and homemade buffer filter paper-based homemade spin column (Lane Q/H), followed tobacco total RNAs purified by using homemade buffer with Qiagen original spin column (Lane H/Q), filter paper recharged used spin column (Lane H/R) and filter paper-based homemade spin column (Lane H/H). The start material amount is 100 mg tobacco young leave in purification experiments using Qiagen spin column, and filter paper recharged used spin column, and 50 mg for homemade spin column purification. All these purifications used 50 μl elution buffer. (E) UV spectrum of tobacco total RNA purified using filter paper recharged spin column with Qiagen RNeasy^®^ plant mini kit buffers (Q/R, blue curve) or homemade buffers (H/R, red curve). Y-axis is UV absorbance, and the X-axis is wavelength. (F) Amplification plots of three duplicated qRT-PCR reactions contain cDNA amounts to 2.5 ng RNA purified using Qiagen kit (Q/Q, Blue curves) and three duplicated qRT-PCR reactions with cDNA amounts to 2.5 ng RNA purified using filter paper recharged spin column with homemade buffer (H/R, Red curves).

UV spectrum meter analysis shown tobacco leaf DNA purified using filter paper-based recharged spin column with either Qiagen kit buffer or homemade buffer all shown standard DNA absorbance curve with the highest peak at 260 nm (Fig 3B).

To further check the quality of purified DNA, we adopted qPCR to see whether DNA purified by commercial kit and filter paper-based spin column with the homemade buffer can be amplified in the same efficiency. As shown in Fig 3C, amplifications of the same amount of DNAs purified by Qiagen kit and filter paper recharged spin column with homemade buffer shown the same amplification plots with nearly identical Cq value across the threshold. Based on assumption that PCR amplicon double their amount after each PCR cycle, there are no difference in PCR analysis for these DNAs (Fig 3C), and confirmed that DNA purified using the filter paper-based with homemade buffer approach is the same quality and has no additional inhibitor of PCR reaction as DNA purified using commercial kit.

DNA purified by using recharged and homemade spin column is more suitable for low throughput experiments, such as confirmation of transgenic plants via PCR. We also successfully use tomato and tobacco DNAs purified by homemade spin column with homemade buffer for molecular marker development experiments, including PCR based Cleaved Amplified Polymorphic Sequences (CAPS) or Kompetitive Allele Specific PCR (KASP) (unpublished data).

In addition to genomic DNA purification, we tested an in-house protocol and buffered described by Yaffe et al [11] for plant RNA purification. Our experiments confirmed that this protocol could be adopted for purification of tobacco plant RNA from leaf tissue using filter paper-based spin column as well. As shown in Fig 3B, the performance in the purification of tobacco total RNA using filter paper-based spin column with homemade buffers is similar to that of Qiagen Plant RNeasy mini kit. RNAs purified by filter paper recharged spin column with homemade buffer show the same pattern and density of EB stain rRNA bands as those purified by Qiagen kit, which indicated the purified RNAs have similar integrity and abundance (Fig 3D).

Analysis using UV spectrum meter shown standard RNA absorbance curve for tobacco leaf RNA purified using filter paper recharged spin column with Qiagen kit buffer or homemade buffer (Fig 3E). qRT-PCR analysis of RNAs purified by these different method also shown no noticeable difference as to the amplifications efficiencies between RNAs purified by Qiagen kit and RNA purified using filter paper bases spin column with homemade buffer, since Cq of these reactions are similar as indicated the cross point of their amplification curves with threshold (Fig 3F). Such results suggested the same efficiency in both of reverse transcription or real time PCR reaction using RNAs purified by these different approaches. Therefore, filter paper based spin column can be adopted for plant RNA followed commercial kit protocol using kit buffer leftover to save resources, or using an in-house protocol with homemade buffer to further reduce the cost in the lab.

RNA purified by filter paper based spin column using a homemade buffer is ready to use for many downstream experiments, such as RT-PCR analysis of transgene expression in a transgenic plant, RACE for cDNA cloning, and even construction of mRNA deep sequencing libraries (unpublished data).

## Discussion

Development of filter paper-based spin column would be useful for the utility of filter paper in nucleic acid purification since the spin column is a more adopted format which can operate using a conventional desktop centrifuge for low throughput bench-scale nucleic acid experiments required in daily practices of the biology lab. Here we found either recharged or homemade filter paper-based spin columns are suitable for such an application, and they can incorporate with commercial kits or in-house protocol by replacing expensive commercial spin column. The apparent advantage of such practice is to save on laboratory monetary resources. For instance, the price per commercial spin column usually ranges from $0.3 to over $1 on the market. In preparing recharged or homemade spin column with filter paper, the filter paper discs cost less than 1 cent per column, though plastic ware of homemade spin column is about $0.1, all plastic wares can be reused by washing using bleach solution which costs very little. Of course, labor in spin column preparation should be considered, but the extra effort should be acceptable for low throughput applications, mainly followed the simplified recharging processes without adding plastic fixing ring and support frit which commonly adopted in many types of commercial spin column.

Another way to save lab resources is to take advantage of the extra buffer of commercial kit or using a less expensive homemade buffer. To use the extra buffer, some labs regenerated spin column after regeneration treatment, such as using MaxXBBond regeneration kit [11]. Here we used filter paper to recharge spin column which is easy to perform. Also, using homemade spin column based on 0.5 ml tube can start from reduced amount sample and buffer, and therefore, more efficient in using the commercial kit buffer leftover. Purified nucleic acid by this way still yield final nucleic acid at μg scale and is enough for many downstream applications. The less expensive homemade buffer would be another attractive choice for reducing the cost. The homemade buffer used in in-house protocol might result in higher yield than commercial kit, and also save more than 75% in total expense compared to commercial kit [10, 11, 16].

We found that filter paper has the weak binding ability to nucleic acids in lysate without adding a binding buffer in the protocol of Qiagen DNeasy or RNeasy plant kit. Therefore, we used filter paper spin column to filter cell debris instead of using QIAshredder spin column from Qiagen kit. This filtering step might reduce final yield about 10% based on experiments using cleared lysate (data not shown), but it could help to increase the amount of cleared lysate for samples which is hard to get clean lysate by centrifugation only. The similar strategy can apply to filter cell debris of *E. coli* for quick plasmid DNA purification (data not shown) as well.

As a solid phase nucleic acid purification approach, the protocols adopted for filter paper-based spin column eliminate the usage of toxic solvent, such as chloroform or phenol, and is safe to conduct. Punching filter paper disc process is safe while punching and cutting silica-based glass fiber filter might release airborne glass fiber which is a potential hazard to operator [20]. Also, recycle used spin column by recharging help to reduce the releasing of plastic waste from the lab to the environment and therefore is environment-friendly.

Apart from these advantages, some issues should be cautious in the application of the filter paper recharged or homemade spin column. One is the centrifuge speed. For the recharged spin column with V shape bottom or homemade spin column based on 0.5 ml tube, they should be centrifuged at relatively lower speed when filling with a solution. We usually centrifuge them no more than 8000 × g. Otherwise, it might result in leaking of the solution which does not pass through the filter and in turn reduce the binding of nucleic acids. However, when centrifuge these spin column without a solution, such as at the step of drying filter, it would be OK to centrifuge at full speed of microcentrifuge (>16000 × g). As to spin using a spin column with the net structure at the bottom to support filter discs (Fig 1A), those spin columns can be centrifuged as the full speed of micro centrifuge all the time. However, in steps of binding plant genomic DNA or RNA, we still prefer relative lower speed, such as no more than 8000 × g during binding and washing step, except the last drying filter paper step, which needs full speed to eliminate residue ethanol.

In the purification of plant DNA, we found that the final DNA elution might contain RNA if samples contain high abundant RNA or use less RNase in lysis buffer. DNA with RNA can result in additional low molecular smear visualized in agarose gel electrophoresis, and an also higher ratio of absorbance at 280/260 in UV spectrometer measurement due to higher 280 value. Although DNA with RNA might not affect PCR, it might lead to overestimated DNA concentration in calculation based on UV absorbance value at 280 nm. In case of this, we recommend reducing the amount of sample, such as using 50 mg per purification experiment or extend the time of lysis. We do not recommend increase RNase amount since the cost of RNase is the major portion of total expense in purification using a homemade buffer. For instance, 4ul RNase from Qiagen will cost more than 30 cents based on marker price. We sometimes use a reduced amount of RNase in plant genomic DNA purification with longer time of incubation or spin longer time at centrifuge. Purified DNA can also be quantified by approaches other than UV spectrometer for accurate concentration for experiments sensitive to DNA input amount. For tobacco, we usually use fluorometer and an H33258 dye-based assay to quantify tobacco genomic DNA for PCR based molecular marker analysis [21]. DNA quantification based on UV absorbance at 280 nm might not be consistent for DNA purified from different plant species using different commercial kits [22].

As for RNA purification, DNA contamination needs to be considered [23]. Though we found that filter paper-based spin column did accommodate with in-column DNA digestion protocol as suggested by Qiagen kit manual, we still suggest treating RNA elution using DNase-free^TM^ kit (Invitrogen) or similar kits to eliminate remaining DNA. This step is important for DNA contamination sensitive experiments, such as qRT-PCR for quantifying of low expressed genes. On another hand, if the abundance of detected transcript sequence is much higher than that of the background of DNA, such as in detection of pathogen transcript [9, 19], the DNA treatment might be ignored for reducing the cost if no reverse transcription control could verify the situation. In experiments like a Rapid amplification of cDNA ends (RACE), which is a PCR using one gene-specific primer combined with one arbitrary primer anneal to sequence of polyT adaptor or 5’ adaptor added to cDNA, contaminated DNA in RNA might not be a big issue because DNA cannot be amplified in the PCR with only one gene specific primer, and no additional DNase treatment step also helps to eliminate the worry of reducing RNA integrity which is important for RACE like experiments.

At last, the successfully nucleic acid purification using filter paper mainly relies on cellulose, the major component of filter paper. Cellulose seems has the similar feature as silica-based material to bind nucleic acids in chaotropic condition, but only secondary fibril-associated cellulose found in filter paper was found able to isolate a wide range of nucleic acids, another type of cellulose with smooth surfaces was found is less efficient in recover DNA from solution [5]. On another hand, we found that the cellulose-based filter paper is more efficient in the purification of high molecular weight genomic DNA, which is in consistent with similar phenomenon reported for Promage’s Paramagnetic cellulose. DNA purified using this commercial format of cellulose product shown much better performance in plant genomic DNA purification [6]. In addition, we found that filter paper is less efficient for plasmid DNA purification followed miniprep protocol. These phenomena suggested that cellulose based material does not exactlly followed the same mechanism as silica-based material in binding and elution process for nucleic acids. Unlike silica-based nucleic acids purification process which has been extensively studied [24], less report associated with interaction between cellulose and nucleic acids [9], and therefore, more studies are needed to facilitate application of the filter paper-based nucleic acid purification.

## Conclusions

We found that filter paper can replace silicon material for purification of many types of nucleic acids. Filter paper can be easily adopted in the form of recharged used spin column or homemade spin column for low throughput application using commercial kit buffer leftover or homemade buffers to reduce the cost, therefore, can be an important component for nucleic acid purification in molecular biology laboratories.

## Acknowledgments

We are thankful to Ann Piotrowski for help in planting the plants in the greenhouse and field.

